# DeepGeni: Deep generalized interpretable autoencoder elucidates gut microbiota for better cancer immunotherapy

**DOI:** 10.1101/2021.05.06.443032

**Authors:** Min Oh, Liqing Zhang

## Abstract

Recent studies revealed that gut microbiota modulates the response to cancer immunotherapy and fecal microbiota transplantation has clinical benefit in melanoma patients during the treatment. Understanding microbiota affecting individual response is crucial to advance precision oncology. However, it is challenging to identify the key microbial taxa with limited data as statistical and machine learning models often lose their generalizability. In this study, DeepGeni, a deep generalized interpretable autoencoder, is proposed to improve the generalizability and interpretability of microbiome profiles by augmenting data and by introducing interpretable links in the autoencoder. DeepGeni-based machine learning classifier outperforms state-of-the-art classifier in the microbiome-driven prediction of responsiveness of melanoma patients treated with immune checkpoint inhibitors. DeepGeni-based machine learning classifier outperforms state-of-the-art classifier in the microbiome-driven responsiveness prediction of melanoma patients treated with immune checkpoint inhibitors. Also, the interpretable links of DeepGeni elucidate the most informative microbiota associated with cancer immunotherapy response.

## Background

Recent studies have found that the composition of the gut microbiome modulates the response to cancer immunotherapies [1-3]. Immune checkpoint inhibitors (ICIs) that block immunosuppressive molecules of tumor cells, thereby inducing host immune response are highly effective for only a subset of patients (∼40%) [4]. The gut microbiome has been reported as a major extrinsic modulator to responses of ICIs such as anti-PD-1. In mice, fecal microbiota transplantation (FMT) from responders to non-responders promotes the efficacy of anti-PD-1 therapy in non-responders [1-3]. More recently, first-in-human clinical trials observed the clinical benefit of responder-derived FMT in melanoma patients [5, 6]. Although a favorable gut microbiome is associated with response to anti-PD-1 therapy, its composition and the specific mechanisms affecting host immune response remain unclear [7].

Determining the key microbiota affecting individual responses to cancer treatment is crucial for advancing precision oncology. However, this is challenging due to the limited available data sets, thereby lack of generalizability in statistical and machine learning models. For example, multiple studies on small melanoma cohorts have reported gut bacteria associated with response to ICI therapy [1, 2, 8-10], but unfortunately, there are discrepancies in the findings [7]. Many bacteria reported by those studies did not appear in multiple studies at the species level except *Faecalibacterium prausnitzii* and *Bacteroides thetaiotaomicron*. Also, previous attempt to train machine learning classifiers on microbiome profiles has shown relatively low accuracy in the prediction of ICI response on unseen data [11]. This suggests the need for curation of massive-scale studies to obtain statistical power to generalize microbial signatures to unseen data.

Nevertheless, recent advances in artificial intelligence, especially deep learning models for domain generalization may hold promise in generalizing microbial signatures. Domain generalization, also called out-of-distribution generalization, aims at learning models that can be generalized to an unseen domain without any foreknowledge [12]. Domain generalization techniques usually require data from multiple domains or sufficient enough to simulate domain shifts, and the limited availability of microbiome data often restricts the application of the techniques. However, more recent studies proposed data augmentation approaches, circumventing the limitation. Especially, DeepBioGen showed promise in augmenting limited sequencing data, including microbiome profiles, and improving the generalizability of classification models.

Well-generalized and accurate deep learning models have the potential to be a key part of clinical decision-making in precision medicine [13, 14]. Despite the remarkable performance, deep learning models are usually black-box and difficult to interpret, which hampers their adoption in clinical practice as clinicians and decision-makers prioritize the explainability of the predictions [15]. Also, interpretable models may provide useful insight into the underlying mechanisms connecting gut microbiome and host immune response.

In this study, DeepGeni, a deep generalized interpretable autoencoder, is proposed to unveil the gut microbiome associated with ICI response (Figure 1). The previous study has shown that a deep autoencoder can produce a highly effective representation of microbiome profiles [16]. Also, a flexible autoencoder model has been developed for interpretable autoencoding without a significant loss of reconstruction accuracy [17]. By augmenting microbiome profiles and by introducing explainable links in the autoencoder, DeepGeni improved the generalizability and interpretability of the learned representation of microbiome profiles. DeepGeni-based classifiers outperform a state-of-the-art classifier in predicting ICI response using microbiome profiles. Also, interpretable links of DeepGeni reveal important taxa for ICI response prediction, and the identified taxa are either associated with prolonged progression-free survival in melanoma patients treated with ICI therapy or differentially abundant between responders and non-responders.

**Figure 1.**
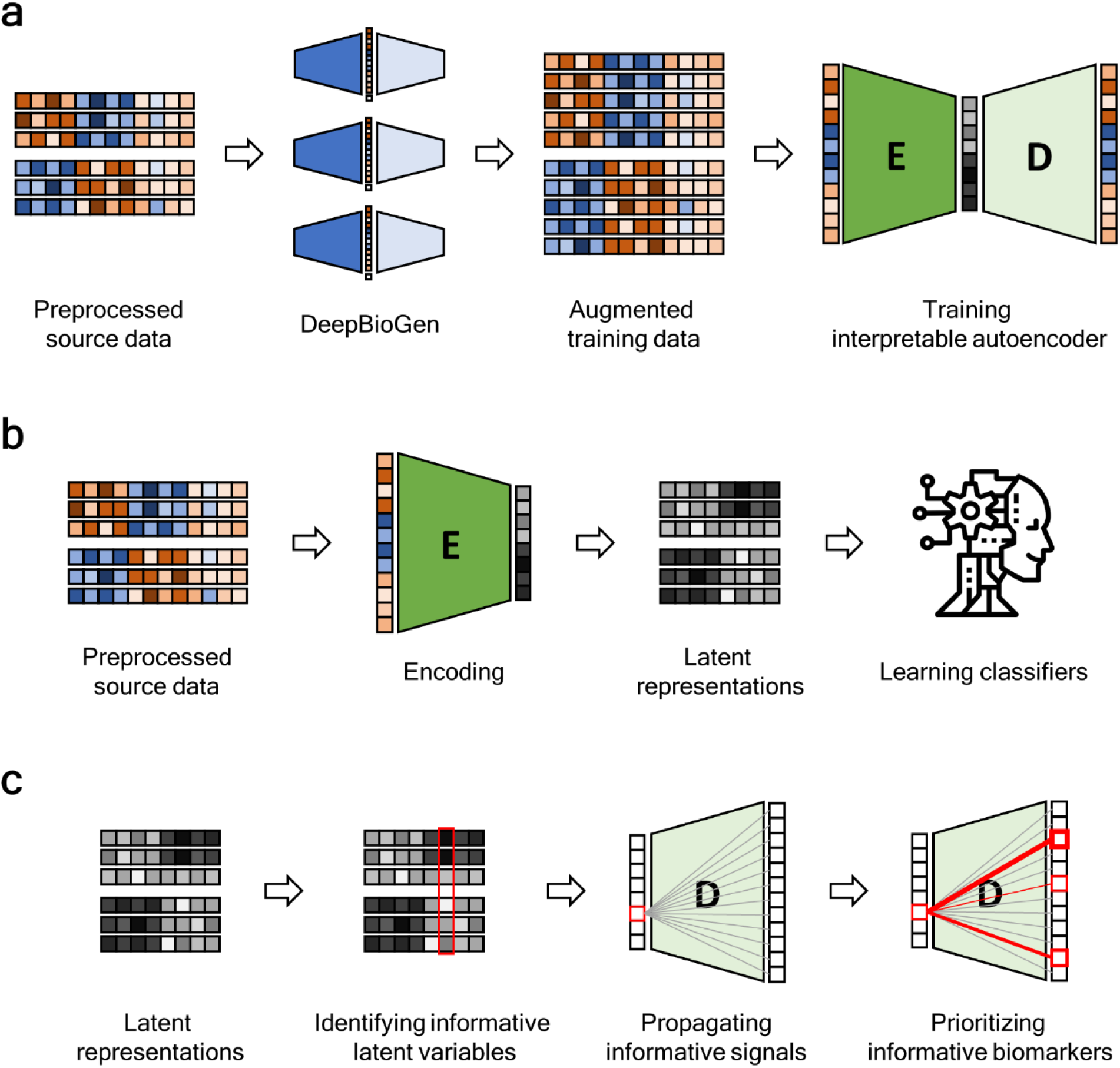
Overview of DeepGeni analysis

## Methods

### Datasets

Gut microbiome data of melanoma patients treated with ICI therapy were collected from four shotgun metagenomic studies [1, 2, 9, 18]. This study focused on samples gathered before ICI therapy and excluded the other samples taken after ICI administration. Patients’ responsiveness to ICI therapy was evaluated with RECIST 1.1 criteria where complete or partial responses are classified as responders and stable or progressive disease states as non-responders [19]. Since Peters et al.’s data did not have an explicit classification of responsiveness, patients with over 6 months of progression-free survival were regarded as responders and the others as non-responders as suggested by Limeta et al. [11]. In total, 130 melanoma patients (66 responders and 64 non-responders) were used (Table 1).

**Table 1.** Summary of gut microbiome datasets derived from shotgun metagenomic sequencing

| Dataset name | # of total samples | # of responders | # of non-responders | Publish ed year | ICI therapy | Reference |
| --- | --- | --- | --- | --- | --- | --- |
| Gopalakrishnan | 25 | 14 | 11 | 2018 | Anti-PD-1 | [1] |
| Matson | 39 | 15 | 24 | 2018 | Anti-PD-1 | [2] |
| Frankel | 39 | 19 | 20 | 2017 | Anti-PD-1, Anti-CTLA-4, and both | [9] |
| Peters | 27 | 18 | 9 | 2019 | Anti-PD-1, Anti-CTLA-4,<br>and both | [18] |

Raw sequencing reads were filtered with FASTP and processed with mOTUs2, a phylogenetic z (mOTU) profiler [20, 21]. Processed microbiome profiles containing read counts for each phylogenetic marker gene and each patient were acquired from Limeta et al. [11]. Read counts were normalized by the total number of reads for each patient, and then log2-transformed. In total, 7,727 mOTUs (features) were considered in an initial input.

### Microbiome profile augmentation with DeepBioGen

DeepGeni utilizes DeepBioGen, a sequencing profile augmentation procedure that generalizes the subsequent trainable models with the augmented data (Figure 1a). Visual patterns of source microbiome profiles are established with feature selection followed by feature-wise clustering.

Wasserstein generative adversarial network (GAN) equipped with convolutional layers capturing the visual patterns generates realistic profiles and augments source data. The augmented training data can enhance the generalizability of the subsequent models such as machine learning classifiers to unseen data. In this study, DeepBioGen parameters were set to default, otherwise, configured following the guideline described in the original paper. Test data has been excluded from any estimation of the parameters. Out of 7,727 mOTU features, 256 features were selected by fitting extremely randomized trees on source data [22]. The number of feature-wise clusters and the number of GAN models were estimated by calculating the within-cluster sum of squared errors in source data with reduced features.

### Generalized autoencoder with interpretable links

Autoencoder consists of encoder and decoder functions that are approximated by neural networks. The encoder maps the input data points into latent space and the decoder reconstructs the input from the mapped latent representations. During training, the autoencoder tries to minimize the gap between the input and the reconstruction by adjusting weights of neural networks based on back-propagated signals from reconstruction loss term. Formally, the reconstruction loss can be written as,

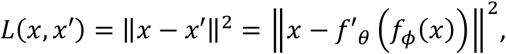

where *x* and *x*′ are the input and the reconstruction, *f*_*ϕ*_(·) and *f*′_*ϕ*_(·) are encoder and decoder functions in which *ϕ* and θ are their weights, respectively. The latent representation usually has a smaller dimension than the original input but it contains concentrated information that can be used to reconstruct the original input with minimal error. Although the latent representation may hold essential information in a condensed form, it is not directly interpretable because of the non-linear relationship between latent and original features.

Svensson et al. suggested a flexible autoencoder model removing non-linearity in decoder function, opening up a possibility to retain interpretability without ruining reconstruction quality [17]. The non-linearity of the autoencoder comes from a non-linear activation function applied to the weighted sum of the preceding inputs. By removing the activation function in the decoder part, direct linear links from the latent layer to the output layer can be obtained. In this study, simple autoencoder architectures composed of three dense layers were utilized: input layer, latent layer, and output layer. The number of nodes of the input and output layer is the same as that of the input. Four different sizes of latent nodes were examined: 128, 64, 32, and 16. The augmented training data consisting of source and augmented data was used to train the autoencoder. After training, the encoder part was used to produce latent representations of the augmented training data. Test data was isolated from any steps of autoencoder training.

### Generalized latent representations for predicting ICI responses

To estimate the usefulness of the latent representations derived from the generalized autoencoder, prediction models classifying ICI responses were built on the representations (Figure 1b). Three machine learning algorithms, support vector machine (SVM), random forest (RF), and multi-layer feedforward neural network (NN) were used to train the models:. Prediction performance was evaluated with two approaches. The first approach, similar to Limeta et al., utilizes the most recent dataset in Peters et al. (Peters) as test data and the remaining data pooled together as source data. The other approach is cross-study validation that iterates over datasets, leaves one dataset as test data, uses the remaining as source data, and averages over results. For both approaches, five-fold cross-validation on the learned representation of source data was conducted to optimize hyper-parameters of the classification algorithms. Hyper-parameter space was explored with grid search and the parameter grid is described in Supplementary Table S1. With the best hyper-parameters, classifiers were trained on representations of the entire source data and evaluated on test data. Area under the receiver operating characteristics curve (AUC) was used to assess the prediction performance.

### Extracting informative microbiota from interpretable autoencoder

To interpret the latent representations that improve the prediction of ICI response, the most informative latent variables were selected based on feature importance estimated by extremely randomized trees [22]. The informative signals of the selected latent variables were propagated through direct links in the decoder network (Figure 1c). Out of 128 latent variables, ten of the most informative variables were considered for further analysis. For each variable, the links were ranked by the absolute value of their weights and, out of 256 links, the top 20 were selected. After the corresponding output nodes connected to the top 20 links were mapped to mOTUs in a one-to-one manner, the specified 20 mOTUs were listed into a set of candidates. By iterating over the ten latent variables, the ten sets of candidates were merged into a unique set of candidates. The whole process was repeated four times by dropping one data set at a time and using the rest for better generalizability. The finalist was acquired by taking the intersection of the four sets of candidates and it contains 14 mOTUs.

### Statistical Analysis

To assess the impact of the identified informative mOTUs on ICI responsiveness, progression-free survival analysis that is a primary endpoint of clinical oncology studies was conducted. Data in Peters et al. (N=27) has progression-free survival and was used in the analysis. For each mOTU, the second quartile (median) was used as a cut-off for high abundance. The Kaplan-Meier plot was drawn and the log-rank test was conducted for statistical significance. Wilcoxon rank-sum test was used to determine differentially abundant taxa.

## Results

### Improved prediction of ICI response with generalized interpretable autoencoder

We evaluated the prediction performance of machine learning classifiers utilizing DeepGeni, a deep generalized interpretable autoencoder. The classifiers were learned to predict a binary class of ICI treatment (responder vs non-responder) based on the latent representation of microbiome profiles. Test data has been excluded from the whole process of generalizing and training the autoencoder of which encoder part produces the latent representation. DeepGeni-based classifiers were compared to classifiers trained on three different settings without augmentation: 1) Initial data of 7,727 mOTU features without feature selection or latent encoding, 2) Feature selected data (256 mOTU features) without latent encoding, 3) Feature selected data with latent encoding. For each approach, out of three classification algorithms (SVM, RF, and NN), the best performing one was selected. Also, a state-of-the-art approach that selects differentially abundant mOTU features and applies a random forest classification algorithm was included in the comparison. As an independent validation setting, the most recent study’s data (Peters) was used as test data and the rest as source data for training classifiers.

Remarkably, DeepGeni-based NN classifier surpasses not only state-of-the-art classifier (Limeta et al.) but the best classifiers of other approaches (Figure 2). In addition, the rest of DeepGeni-based classifiers (SVM and RF) show better performance than the classifiers of other approaches (Table S2). Also, DeepGeni-based SVM classifier outperforms other classifiers in the cross-study validation setting, displaying the highest generalizability across different studies (Table 1).

**Figure 2.**
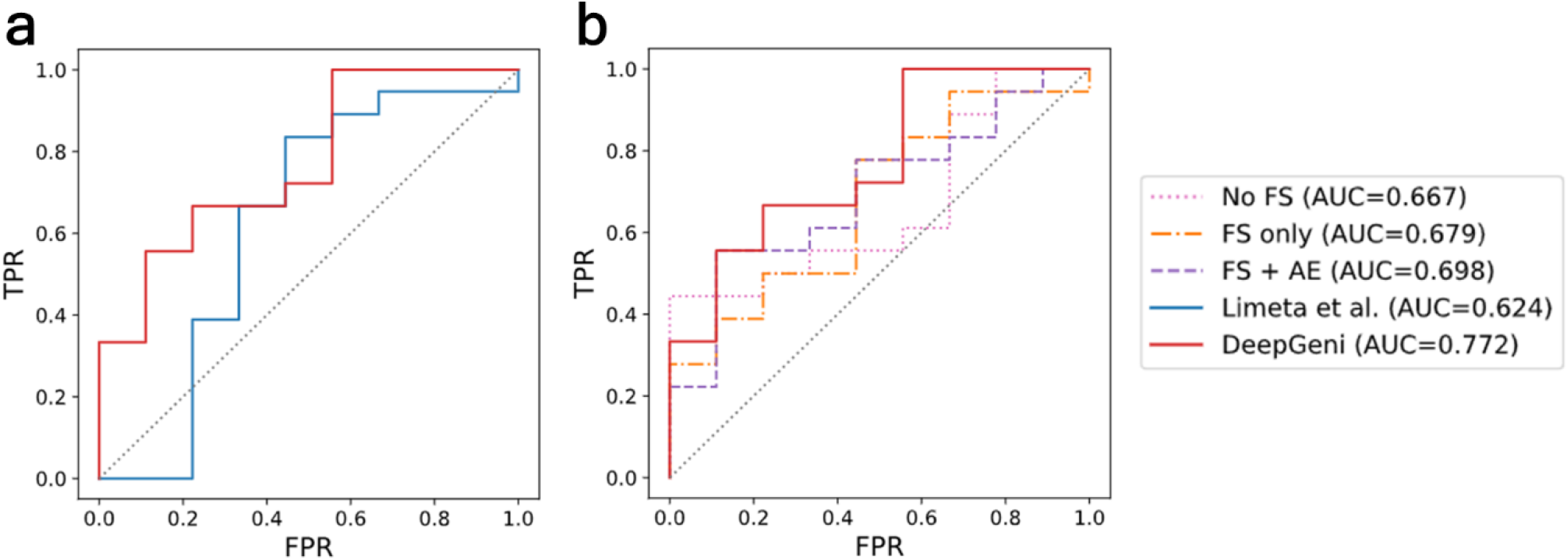
Receiver operating characteristics (ROC) curves of the best classifier for each method

**Table 1.** Averaged AUC in cross-study validation setting

| Approach | No FS |  |  | FS only |  |  | FS + AE |  |  | DeepGeni<br>(FS + DBG + AE) |  |  |
| --- | --- | --- | --- | --- | --- | --- | --- | --- | --- | --- | --- | --- |
| Algorithm | SVM | RF | NN | SVM | RF | NN | SVM | RF | NN | SVM | RF | NN |
| AUC | 0.52 | 0.522 | **0.556** | 0.564 | 0.551 | **0.585** | **0.602** | 0.57 | 0.598 | **0.626** | 0.579 | 0.609 |
| STD | 0.156 | 0.074 | 0.07 | 0.107 | 0.103 | 0.08 | 0.06 | 0.053 | 0.045 | 0.209 | 0.09 | 0.221 |

- FS: feature selection; AE: autoencoder; DBG: DeepBioGen

### Key microbiota relevant to ICI response extracted from generalized interpretable autoencoder

The finalist of ICI-response-relevant key microbiota was identified by propagating informative signals through the interpretable links from latent variables that play a major role in inducing the superior ICI response prediction. The finalist consisting of 14 mOTUs categorized into seven families were validated with the literature and statistical tests. Previous studies have reported 12 of the 14 generally in the broader taxonomic categories. However, the finalist generally shows a higher resolution of microbiota associated with ICI therapy in taxonomic identification (Table 2). Interestingly, two ICI-therapy-relevant gut bacteria, *Eggerthella lenta* and unknown *Lactobacillales*, were not reported in previous studies, thus providing new microbe markers for future studies. It is worth noting that the genus Subdoligranulum is closely related to the Faecalibacterium genus. Furthermore, five species, including *Lactobacillus plantarum*, unknown *Ruminococcaceae*, and three unknown *Clostridiales*, displayed statistical significance in differentially abundant testing (unadjusted, Wilcoxon’s rank-sum test). Besides, a high abundance of unknown *Eubacterium* species was significantly associated with prolonged progression-free survival in ICI-treated melanoma patients (Figure 3).

**Table 2.** The finalist of ICI-response-relevant key microbiota

| mOTU_v2 ID | Consensus taxonomy | Order | Family | Genus | Specified level | Prev level | H-Res | P-val |
| --- | --- | --- | --- | --- | --- | --- | --- | --- |
| ref_mOTU_v2_0036 | Enterobacteriaceae sp. | Enterobacteriales | Enterobacteriaceae | Escherichia /Shigella | Species | Species [2] | - |  |
| ref_mOTU_v2_0154 | Lactobacillus plantarum | Lactobacillales | Lactobacillaceae | Lactobacillus | Species | Family [2] | Yes | * |
| meta_mOTU_v2_6288 | unknown Lactobacillales |  | unknown Lactobacillales | unknown | Family | - | - |  |
| ref_mOTU_v2_0642 | Eggerthella lenta | Eggerthellales | Eggerthellaceae | Eggerthella | Species | - | - |  |
| ref_mOTU_v2_0884 | Anaerotruncus colihominis | Clostridiales | Ruminococcaceae | Anaerotruncus | Species | Family [1] | Yes |  |
| ref_mOTU_v2_4738 | Subdoligranulum sp. |  |  | Subdoligranulum | Species | Family [1] | Yes |  |
| ref_mOTU_v2_0281 | Ruminococcus lactaris |  |  | Ruminococcus | Species | Genus [1, 8] | Yes |  |
| meta_mOTU_v2_6557 | unknown Ruminococcaceae |  |  | unknown Ruminococcaceae | Genus | Family [1] | Yes | ** |
| meta_mOTU_v2_6657 | unknown Eubacterium |  | Eubacteriaceae | Eubacterium | Species | Genus [1, 8] | Yes | # |
| meta_mOTU_v2_5411 | unknown Clostridiales |  | unknown Clostridiales | unknown | Family | Order [1] | Yes |  |
| meta_mOTU_v2_5669 | unknown Clostridiales |  |  |  |  |  |  | * |
| meta_mOTU_v2_6760 | unknown Clostridiales |  |  |  |  |  |  | * |
| meta_mOTU_v2_6795 | unknown Clostridiales |  |  |  |  |  |  | * |
| meta_mOTU_v2_7550 | unknown Clostridiales |  |  |  |  |  |  |  |

- \*: $p < 0.05$ , Wilcoxon's rank-sum test on differential abundance; \*\*: $p < 0.01$ , Wilcoxon's rank-sum test; #: $p < 0.05$ , log-rank test on progression-free survival distribution difference; H-Res indicates whether the specified taxonomic level is in higher resolution than the previously specified level in other studies.

**Figure 3.**
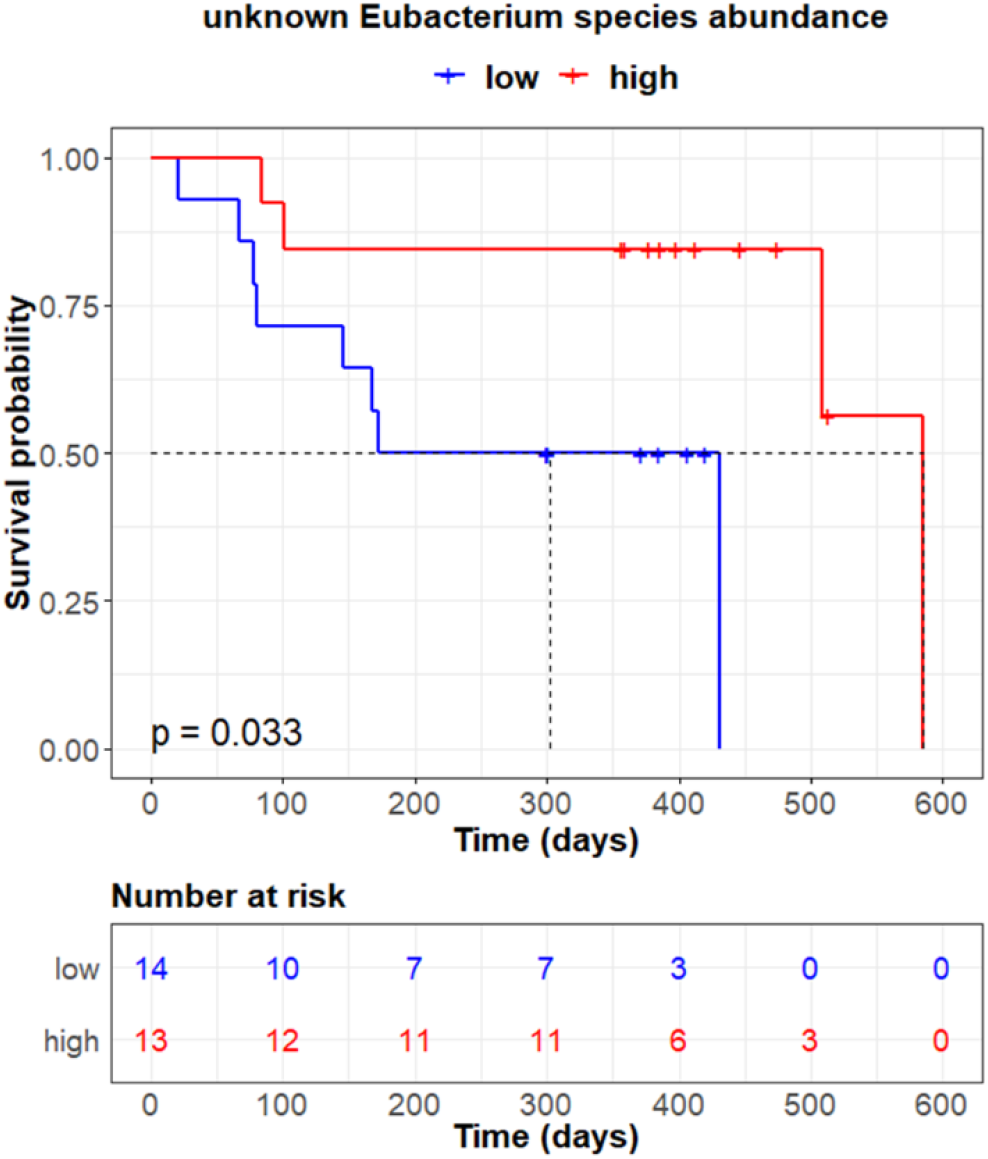
Kaplan-Meier plot of progression-free survival by relative abundance of unknown Eubacterium species

## Discussion

DeepGeni is a generalized interpretable autoencoder not only boosting ICI response prediction accuracy in an independent study but providing interpretable links to identify informative taxa contributory to modulate ICI response. The improved generalizability of DeepGeni is supposed to be derived from augmented microbiome data generated by DeepBioGen, a GAN-based data augmentation procedure. The latent representation learned by the generalized autoencoder with the augmented data can enable to train classifiers more resilient to unseen data distributions. Also, DeepGeni extracted microbial species informative to predict ICI response in higher resolution than other studies. The specified species could be a helpful basis for establishing ICI-promoting FMT guidelines to specify donor and donee. Moreover, the identified species may offer a possibility to develop pre or probiotics targeting improved outcomes of ICI therapy.

Although this study produces the generalized list of ICI-response-relevant key microbial taxa over the available datasets, the ability to statistically validate the identified microbial taxa is bounded by the size of the available data. This could limit the possibility of being validated for some of the key microbial taxa as they were identified by taking advantage of the out-of-distribution augmented data and it may not be eligible to use the augmented data for statistical validation. However, there still remains the possibility of being validated in larger data sets once they become available.

DeepGeni was applied to examine microbiome modulating ICI response in this study but it is highly extensible for identifying microbiome-driven human phenotypes or even for applying other types of biological and ecological data such as genome and metagenome profiles.

## Conclusion

We proposed DeepGeni, a generalized interpretable autoencoder that learns a latent representation of microbiome profiles. The learned representation can improve ICI response prediction on unseen data and suggest the most informative microbial taxa involved in modulating ICI response.

## List of abbreviations

ICI: Immune checkpoint inhibitor
FMT: fecal microbiota transplantation
mOTU: marker gene-based operational taxonomic unit
GAN: generative adversarial network
SVM: support vector machine
RF: random forest
NN: feedforward neural network
AUC: Area under the receiver operating characteristics curve
ROC: Receiver operating characteristics

## Acknowledgments

This work is partially supported by the funding from Data and Decisions Destination Area at Virginia Tech.

## Author contributions

MO designed the study, collected data, implemented the software, and performed experiments. MO and LZ interpreted the results and wrote the manuscript. All authors read and approved the final manuscript.

## Reference

1. Gopalakrishnan V, Spencer CN, Nezi L, Reuben A, Andrews M, Karpinets T, Prieto P, Vicente D, Hoffman K, Wei S: Gut microbiome modulates response to anti–PD-1 immunotherapy in melanoma patients. Science 2018, 359:97–103.

2. Matson V, Fessler J, Bao R, Chongsuwat T, Zha Y, Alegre M-L, Luke JJ, Gajewski TF: The commensal microbiome is associated with anti–PD-1 efficacy in metastatic melanoma patients. Science 2018, 359:104–108.

3. Routy B, Le Chatelier E, Derosa L, Duong CP, Alou MT, Daillère R, Fluckiger A, Messaoudene M, Rauber C, Roberti MP: Gut microbiome influences efficacy of PD-1–based immunotherapy against epithelial tumors. Science 2018, 359:91–97.

4. Marcus L, Lemery SJ, Keegan P, Pazdur R: FDA approval summary: pembrolizumab for the treatment of microsatellite instability-high solid tumors. Clinical Cancer Research 2019, 25:3753–3758.

5. Baruch EN, Youngster I, Ben-Betzalel G, Ortenberg R, Lahat A, Katz L, Adler K, Dick-Necula D, Raskin S, Bloch N: Fecal microbiota transplant promotes response in immunotherapy-refractory melanoma patients. Science 2021, 371:602–609.

6. Davar D, Dzutsev AK, McCulloch JA, Rodrigues RR, Chauvin J-M, Morrison RM, Deblasio RN, Menna C, Ding Q, Pagliano O: Fecal microbiota transplant overcomes resistance to anti–PD-1 therapy in melanoma patients. Science 2021, 371:595–602.

7. Shaikh FY, Gills JJ, Sears CL: Impact of the microbiome on checkpoint inhibitor treatment in patients with non-small cell lung cancer and melanoma. EBioMedicine 2019, 48:642–647.

8. Chaput N, Lepage P, Coutzac C, Soularue E, Le Roux K, Monot C, Boselli L, Routier E, Cassard L, Collins M: Baseline gut microbiota predicts clinical response and colitis in metastatic melanoma patients treated with ipilimumab. Annals of Oncology 2017, 28:1368–1379.

9. Frankel AE, Coughlin LA, Kim J, Froehlich TW, Xie Y, Frenkel EP, Koh AY: Metagenomic shotgun sequencing and unbiased metabolomic profiling identify specific human gut microbiota and metabolites associated with immune checkpoint therapy efficacy in melanoma patients. Neoplasia 2017, 19:848–855.

10. Vétizou M, Pitt JM, Daillère R, Lepage P, Waldschmitt N, Flament C, Rusakiewicz S, Routy B, Roberti MP, Duong CP: Anticancer immunotherapy by CTLA-4 blockade relies on the gut microbiota. Science 2015, 350:1079–1084.

11. Limeta A, Ji B, Levin M, Gatto F, Nielsen J: Meta-analysis of the gut microbiota in predicting response to cancer immunotherapy in metastatic melanoma. JCI insight 2020, 5.

12. Wang J, Lan C, Liu C, Ouyang Y, Qin T: Generalizing to Unseen Domains: A Survey on Domain Generalization. arXiv preprint arXiv:210303097 2021.

13. Cammarota G, Ianiro G, Ahern A, Carbone C, Temko A, Claesson MJ, Gasbarrini A, Tortora G: Gut microbiome, big data and machine learning to promote precision medicine for cancer. Nature Reviews Gastroenterology & Hepatology 2020, 17:635–648.

14. Wilkinson J, Arnold KF, Murray EJ, van Smeden M, Carr K, Sippy R, de Kamps M, Beam A, Konigorski S, Lippert C: Time to reality check the promises of machine learning-powered precision medicine. The Lancet Digital Health 2020.

15. Wang F, Kaushal R, Khullar D: Should health care demand interpretable artificial intelligence or accept “black box” medicine? : American College of Physicians; 2020.

16. Oh M, Zhang L: DeepMicro: deep representation learning for disease prediction based on microbiome data. Scientific reports 2020, 10:1–9.

17. Svensson V, Gayoso A, Yosef N, Pachter L: Interpretable factor models of single-cell RNA-seq via variational autoencoders. Bioinformatics 2020, 36:3418–3421.

18. Peters BA, Wilson M, Moran U, Pavlick A, Izsak A, Wechter T, Weber JS, Osman I, Ahn J: Relating the gut metagenome and metatranscriptome to immunotherapy responses in melanoma patients. Genome medicine 2019, 11:1–14.

19. Eisenhauer EA, Therasse P, Bogaerts J, Schwartz LH, Sargent D, Ford R, Dancey J, Arbuck S, Gwyther S, Mooney M: New response evaluation criteria in solid tumours: revised RECIST guideline (version 1.1). European journal of cancer 2009, 45:228–247.

20. Chen S, Zhou Y, Chen Y, Gu J: fastp: an ultra-fast all-in-one FASTQ preprocessor. Bioinformatics 2018, 34:i884–i890.

21. Milanese A, Mende DR, Paoli L, Salazar G, Ruscheweyh H-J, Cuenca M, Hingamp P, Alves R, Costea PI, Coelho LP: Microbial abundance, activity and population genomic profiling with mOTUs2. Nature communications 2019, 10:1–11.

22. Geurts P, Ernst D, Wehenkel L: Extremely randomized trees. Machine learning 2006, 63:3–42.

